# Local structural connectivity directs seizure spread in focal epilepsy

**DOI:** 10.1101/406793

**Authors:** Preya Shah, Arian Ashourvan, Fadi Mikhail, Adam Pines, Lohith Kini, Russell T. Shinohara, Danielle S. Bassett, Brian Litt, Kathryn A. Davis

## Abstract

How does the human brain’s structural scaffold give rise to its intricate functional dynamics? This is a central challenge in translational neuroscience, particularly in epilepsy, a disorder that affects over 50 million people worldwide. Treatment for medication-resistant focal epilepsy is often structural – through surgery, devices or focal laser ablation – but structural targets, particularly in patients without clear lesions, are largely based on functional mapping via intracranial EEG (iEEG). Unfortunately, the relationship between structural and functional connectivity in the seizing brain is poorly understood. In this study, we quantify structure-function coupling, specifically between white matter connections and iEEG, across preictal and ictal periods in 45 seizures from 9 patients with unilateral drug-resistant focal epilepsy. We use High Angular Resolution Diffusion Imaging (HARDI) tractography to construct structural connectivity networks and correlate these networks with time-varying broadband and frequency-specific functional networks derived from coregistered iEEG. Across all frequency bands, we find significant increases in structure-function coupling from preictal to ictal periods. We demonstrate that short-range structural connections are primarily responsible for this increase in coupling. Finally, we find that spatiotemporal patterns of structure-function coupling are stereotyped, and a function of each patient’s individual anatomy. These results suggest that seizures harness the underlying structural connectome as they propagate. Our findings suggest that the relationship between structural and functional connectivity in epilepsy may inform current and new therapies to map and alter seizure spread, and pave the way for better-targeted, patient-specific interventions.

## 1. Introduction

Epilepsy is a neurological disorder characterized by recurrent, unprovoked seizures. It affects over 50 million people worldwide [World Health Organization, 2018] and will afflict approximately 1 in 26 people during their lifetime [Hesdorffer et al., 2011]. The most common subtype is *focal* or localization-related epilepsy, in which seizures arise from a specific region in the brain [French, 2007]. Patients with localization-related epilepsy often experience uncontrolled seizures despite medication, leading to neurological and psychiatric co-morbidities, deterioration in quality of life, and up to an eleven-fold increase in mortality rate [Fazel et al., 2013; Kwan et al., 2011].

Recent evidence shows that seizures most commonly arise from abnormal brain networks rather than isolated focal lesions [Bernhardt et al., 2013; Kramer and Cash, 2012]. As a result, researchers are applying graph theoretical methods from the rapidly growing field of network neuroscience to identify brain network abnormalities in epilepsy, in the hope of finding targets for therapeutic interventions. In this approach, investigators map whole-brain structural and functional networks, or “connectomes”, by characterizing connectivity between brain regions based on multi-modal neuroimaging data [Bassett and Sporns, 2017; Bullmore and Sporns, 2009; Rubinov and Sporns, 2010]. Structural brain networks are most commonly derived from diffusion tensor imaging (DTI) tractography [Hagmann et al., 2008]. Functional brain networks are most commonly derived from correlations in signal fluctuations across multiple recording sites from modalities such as resting state functional MRI (fMRI) [Biswal et al., 1995; Salvador et al., 2004], magnetoencephalography (MEG) [Stam, 2004], and electroencephalography (EEG) [Micheloyannis et al., 2006]. These approaches reveal a wide variety of network disruptions in epilepsy patients, both structurally [Raj et al., 2010; Taylor et al., 2015; Vaessen et al., 2012] and functionally [de Campos et al., 2016; Pedersen et al., 2015; Pittau et al., 2012]. While still nascent, this work shows promise for clinical applications, as network-based measures may serve as biomarkers for predicting seizure onset and spread [Burns et al., 2014; Jirsa et al., 2017; Khambhati et al., 2016; Proix et al., 2016], cognitive impairments [Vaessen et al., 2012; Vlooswijk et al., 2011], and outcome following surgical therapy [Goodfellow et al., 2016; Lopes et al., 2018; Sinha et al., 2017].

Most studies of epileptic networks focus solely on either structural or functional connectivity. However, it is commonly understood that the two are tightly linked. In fact, there is great interest in the neuroscience community in elucidating the relationship between brain structure and function. Recent evidence shows that structural and functional brain networks are correlated at multiple temporal and spatial scales, that structural connectivity constrains functional connectivity, and that functional connectivity can modulate structural connectivity via mechanisms of plasticity [Chu et al., 2015; Finger et al., 2016; Greicius et al., 2009; Hagmann et al., 2010; Hermundstad et al., 2013; Hermundstad et al., 2014; van den Heuvel et al., 2009; Honey et al., 2009; Rubinov et al., 2009; Skudlarski et al., 2008; Zhang et al., 2010].

Given the robust coupling between structure and function in healthy brains, disruptions in structure-function coupling can serve as biomarkers of neurological disease, including in epilepsy. For example, Zhang et al. (2011) report that the degree of coupling between resting state fMRI networks and DTI tractography networks is lower in idiopathic generalized epilepsy patients compared with healthy controls, and is negatively correlated with epilepsy duration. Using a similar approach, Chiang et al. (2015) report decreased structure-function coupling in patients with left temporal lobe epilepsy compared with healthy subjects. These two studies employ resting-state fMRI, which characterizes the static, interictal functional epileptic network. However, little is known about the correlation between structural and functional connectivity during seizures. How does structure-function coupling change over the course of seizure evolution? And which particular connections drive these changes? Clinically, it is well understood that focal seizures often quickly spread to distant brain regions, but the relationship of this spread to underlying structure has not been quantified. Understanding where seizures are generated and how they spread has been hampered by sparsely sampled intracranial EEG and lesion-negative clinical brain images, and yet remains vital for planning surgical treatments for epilepsy.

In order to address these questions, we study structure-function coupling in 45 seizures from 9 drug-resistant localization-related epilepsy patients undergoing routine evaluation for epilepsy surgery. To construct time-varying functional connectivity (FC) networks, we utilize clinical recordings from intracranial EEG (iEEG), an invasive method that captures electrical activity from the brain in the form of aggregate local field potentials, at high spatial and temporal resolution [Lachaux et al., 2003; Penfield and Jasper, 1954]. To construct structural connectivity (SC) networks, we analyze High Angular Resolution Diffusion Imaging (HARDI), an advanced diffusion imaging method that can produce robust tractography results in regions of crossing white matter pathways [Tuch et al., 2002]. Below we characterize relationships between these two modalities across time, frequency, and space. We hypothesize that there would be an increase in structure-function coupling during the progression from preictal to ictal states, as seizures spread along structural pathways. Our findings shed light on the pathophysiological processes involved in seizure dynamics, which can ultimately inform new approaches for clinical intervention. We detail these investigations below.

## 2. Materials and Methods

### 2.1 Subjects

We studied nine patients undergoing pre-surgical evaluation for drug-resistant epilepsy at the Hospital of the University of Pennsylvania. Seizure localization was determined via comprehensive clinical evaluation, which included multimodal imaging, scalp and intracranial video-EEG monitoring, and neuropsychological testing. This study was approved by the Institutional Review Board of the University of Pennsylvania, and all subjects provided written informed consent prior to participating.

### 2.2 Intracranial EEG acquisition

Cortical surface and depth electrodes were implanted in patients based on clinical necessity. Electrode configurations (Ad Tech Medical Instruments, Racine, WI) consisted of linear cortical strips and two-dimensional cortical grid arrays (2.3 mm diameter with 10 mm inter-contact spacing), and linear depths (1.1 mm diameter with 10 mm inter-contact spacing). Continuous iEEG signals were obtained for the duration of each patient’s stay in the epilepsy monitoring unit. Signals were recorded at 500 Hz. For each clinically identified seizure event, a board-certified epileptologist precisely annotated the onset time, termination time, seizure type, and electrodes recording artifact signals. Seizure onset times were defined by the earliest electrographic change (EEC) [Litt et al., 2001]. Seizure types were classified using ILAE 2017 criteria [Fisher et al., 2017] as focal aware (previously known as simple partial), focal impaired awareness (previously known as complex partial), or focal to bilateral tonic-clonic (previously known as complex partial with secondary generalization). Furthermore, the onset time of bilateral spread was noted for focal to bilateral tonic-clonic seizures. All annotations were verified and consistent with detailed clinical documentation. To ensure consistency and validity of the captured seizures, we discarded seizures that contained substantial artifacts in all electrodes, events that were very short (< 15 seconds), or those that occurred during sleep. De-identified iEEG recordings are available online on the International Epilepsy Electrophysiology Portal (www.ieeg.org, IEEG Portal) [Kini et al., 2016; Wagenaar et al., 2013].

### 2.3 Image acquisition

Prior to electrode implantation, MRI data were collected on a 3T Siemens Magnetom Trio scanner (Siemens, Erlangen, Germany) using a 32-channel phased-array head coil. High-resolution anatomical images were acquired using a magnetization prepared rapid gradient-echo (MPRAGE) T1-weighted sequence (TR = 1810 ms, TE = 3.51 ms, flip angle = 9°, field of view = 240 mm, resolution = 0.94 × 0.94 × 1.0 mm^3^). High angular resolution diffusion imaging (HARDI) was acquired with a single-shot EPI multishell diffusion-weighted imaging sequence (116 diffusion sampling directions, *b*-values of 0, 300, 700 and 2000 s/mm^2^, resolution = 2.5 × 2.5 × 2.5 mm^3^ resolution, field of view = 240 mm). Following electrode implantation, spiral CT images (Siemens, Erlangen, Germany) were obtained clinically for the purposes of electrode localization. Both bone and tissue windows were obtained (120 KV, 300 mA, axial slice thickness = 1.0 mm).

### 2.4 Region of interest selection

A brain network consists of nodes representing regions of interest (ROIs) within the brain, and edges representing the strength of connectivity between these ROIs. In order to carry out direct quantitative comparisons of structural and functional networks, it was necessary to establish a one-to-one correspondence between functional network nodes and structural network nodes. We therefore determined the location of each electrode in Montreal Neurological Institute (MNI) space and assigned each electrode to its nearest structural region of interest (ROI). Structural ROIs were defined by an upsampled version of the Automated Anatomical Labeling Atlas [Desikan et al., 2006; Tzourio-Mazoyer et al., 2002], which consisted of 600 roughly equally sized (ROI sizes averaging 2.14 +/- 0.28 cm^3^) anatomically constrained regions covering the entire brain with the exception of the cerebellum. We chose this atlas (AAL-600) because it has ROIs of the same order of resolution as iEEG, obeys gross anatomical boundaries, and has successfully been used in prior studies to evaluate structural and functional connectivity patterns in the brain [Hermundstad et al., 2013; Hermundstad et al., 2014].

To determine electrode MNI coordinates, electrodes were first identified via thresholding of the CT image and labeled using a semi-automated process. Each patient’s CT and T1-weighted MRI images were aligned using 3D rigid affine registration, with mutual information as the similarity metric. The T1-weighted MRI images were then aligned to the standard MNI brain using diffeomorphic registration with the symmetric normalization (SyN) method [Avants et al., 2008]. The resulting transformations were used to warp the coordinates of the electrode centroids into MNI space. Co-registrations and transformations were carried out using Advanced Normalization Tools (ANTS) software [Avants et al., 2009; Avants et al., 2011], and the accuracy of each step was confirmed via visual inspection. In our final framework, electrodes served as nodes of the functional networks and the associated structural ROIs served as nodes of the structural networks.

### 2.5 Structural network generation

Diffusion-weighted images were skull-stripped via the FSL brain extraction tool and underwent eddy current and motion correction via the FSL eddy tool [Andersson and Sotiropoulos, 2016]. Next, DWI susceptibility distortions were mitigated using the structural T1-weighted image as follows: T1-weighted images were registered to the b0 image from the DWI scans using FSL FLIRT boundary-based registration [Greve and Fischl, 2009], T1-weighted images were contrast inverted and intensity matched to the DWI image, and finally the DWI scans underwent nonlinear transformation to the T1-weighted scan [Wang et al., 2017]. Following these preprocessing steps, DSI-Studio (http://dsi-studio.labsolver.org) was used to reconstruct the orientation density functions (ODFs) within each voxel using generalized q-sample imaging with a diffusion sampling length ratio of 1.25 [Yeh et al., 2010]. Deterministic whole-brain fiber tracking was performed using an angular threshold of 35 degrees, step size of 1 mm, and quantitative anisotropy threshold based on Otsu’s threshold [Otsu, 1979]. The fiber trajectories were smoothed by averaging the propagation direction with 20% of the previous direction. Tracks with length shorter than 10 mm or longer than 400 mm were discarded, and a total of 1,000,000 tracts were generated per brain. Deterministic tractography was chosen based upon prior work indicating that deterministic tractography generates fewer false positive connections than probabilistic approaches, and that network-based estimations are substantially less accurate when false positives are introduced into the network compared with false negatives [Zalesky et al., 2010].

Subject-level AAL-600 atlases were generated in DWI space by applying the previously generated registration transformations from MNI to T1-weighted space and from T1-weighted space to DWI space. Finally, structural networks were generated by computing the number of streamlines connecting each pair of structural ROIs identified in Section *2.4*. The distribution of mean streamline lengths between each pair of structural ROIs for each patient is illustrated in **Supp. Figure 1**. Streamline counts were subsequently log-transformed to improve normality of the distribution, as is common in prior studies [Bonilha et al., 2015; Park et al., 2017; Taylor et al., 2018; Wirsich et al., 2016].

**Figure 1:**
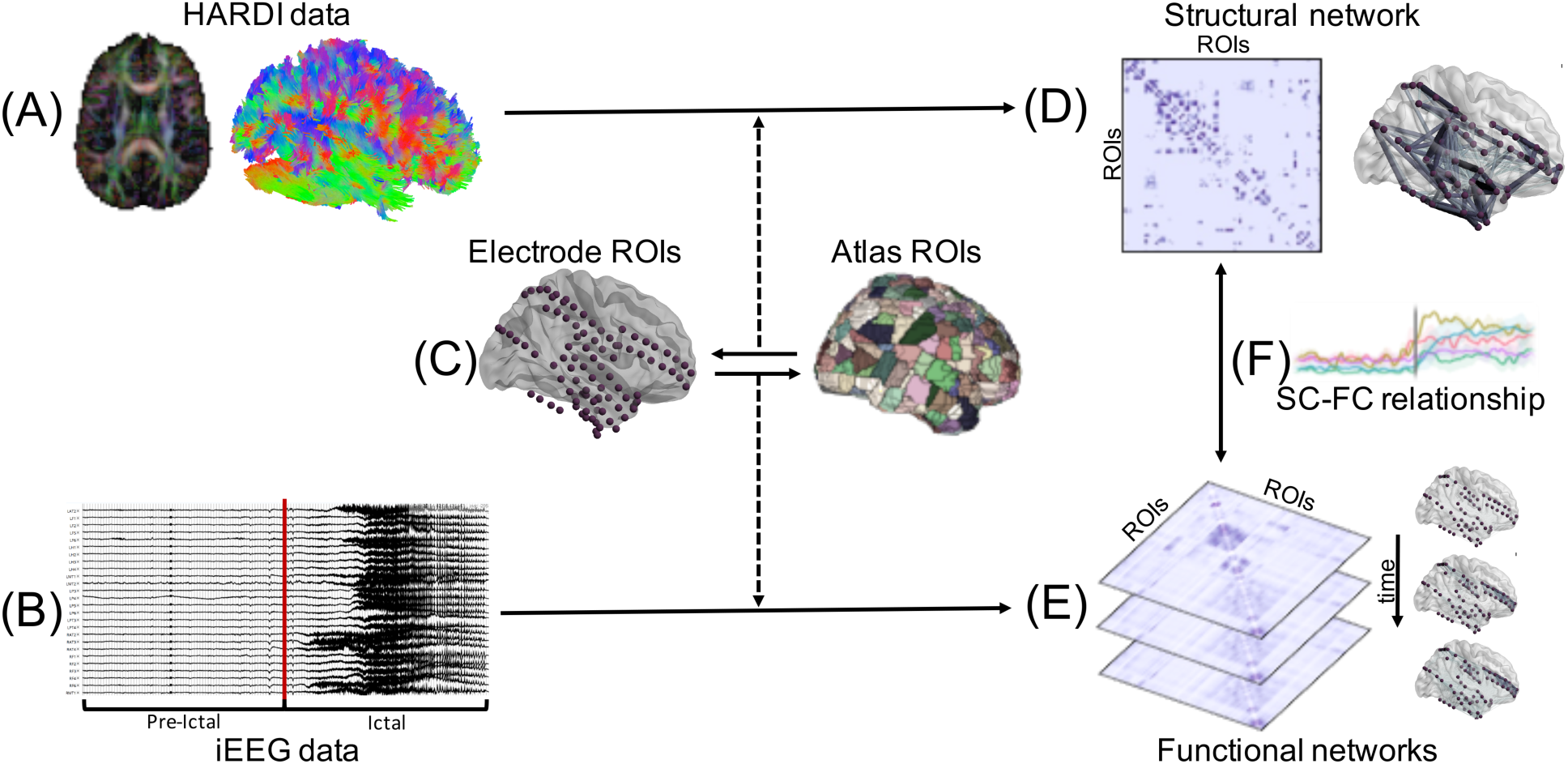
Summary of patient-level SC-FC analysis pipeline. **(A)** HARDI pre-processing and whole-brain tractography was carried out. **(B)** iEEG data were pre-processed and seizures were annotated, with each seizure event consisting of an ictal period and an associated preictal period of equivalent duration. **(C)** Regions of interest (ROIs) were selected via a one-to-one spatial correspondence between electrode centroids and atlas regions. **(D)** The structural connectivity (SC) network was generated using log-normalized streamline counts between atlas ROIs associated with each electrode location. **(E)** Time-varying broadband functional connectivity (FC) networks were generated for each 1s time window by computing correlation between iEEG signals across electrode pairs. Frequency-specific FC networks were similarly computed using coherence between iEEG signals across electrode pairs. **(F)** SC-FC relationships were quantified across time, frequency, and space (see **Methods** for details).

### 2.5 Functional network generation

Each seizure event consisted of an ictal period spanning the time between seizure onset (EEC) and termination, and an associated preictal period of equivalent duration immediately prior to seizure onset. Following removal of artifact-ridden electrodes, intracranial EEG signals for each seizure event were common-average referenced to reduce potential sources of correlated noise [Ludwig et al., 2009]. Next, each event was divided into 1 s non-overlapping time windows in accordance with previous studies [Khambhati et al., 2015; Khambhati et al., 2016; Khambhati et al., 2017; Kramer et al., 2010].

To generate a functional network representing broadband functional interactions between iEEG signals for each 1 s time window, we carried out a method described in detail previously [Khambhati et al., 2017]. Namely, signals were notch-filtered at 60 Hz to remove power line noise, low-pass and high-pass filtered at 115 Hz and 5 Hz to account for noise and drift, and pre-whitened using a first-order autoregressive model to account for slow dynamics. Functional networks were then generated by applying a normalized cross-correlation function *ρ* between the signals of each pair of electrodes within each time window, using the formula:

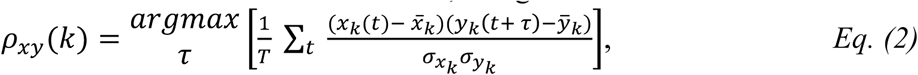

where *x* and *y* are signals from two electrodes, *k* is the 1 s time window, *t* is one of the *T* samples during the time window, and *τ* is the time lag between signals, with a maximum lag of 250 ms. Next, to gain an understanding of the frequency dependence of SC-FC relationships, we generated functional networks across physiologically relevant frequency bands as described in detail in a previous study [Khambhati et al., 2016]. Specifically, multitaper coherence estimation (time-bandwidth product of 5, 8 tapers) was used to compute functional coherence networks for each 1 s window across four frequency bands: α/θ (5-15 Hz), β (15-25 Hz), low-γ (30-40 Hz), and high-γ (95-105 Hz). Both broadband and frequency-specific networks were represented as full-weighted adjacency matrices for each 1 s window in each seizure event.

### 2.6 Structure-function coupling analysis

To quantify the relationship between structure and function in the epileptic brain, we computed the Pearson correlation coefficient between the edges of each SC network and the edges of each broadband FC network, followed by Fisher *r-z* transformation for variance stabilization [Fisher, 1921]. This led to a time series of SC-FC correlations for each seizure event in each subject. To better understand the frequency-dependence of SC-FC coupling, we repeated the same analysis using the frequency-specific functional networks.

Next, to understand the extent to which the resulting SC-FC time series evolve similarly within each subject, we computed the Euclidean distances between these time series for all pairs of seizure events. Importantly, we first time-normalized the time series for each seizure event to span 200 evenly spaced time bins (100 preictal and 100 ictal). Next, for each seizure event, we generated a single vector consisting of the SC-FC time series for all six frequency bands: broadband, α/θ, β, low-γ, and high-γ. Euclidean distances were then computed between all pairs of vectors, comprised of pairs belonging to the same patient and pairs belonging to different patients.

Finally, we wished to assess which edges in the structural network were responsible for the changes in SC-FC correlation between preictal and ictal periods. We therefore first computed a mean ictal and mean preictal broadband functional network for each subject by averaging across seizures events and across windows within each time period. Next, we carried out a virtual edge resection approach, in which we removed an edge from the network and computed the change in SC-FC correlation, Δ*z*(*i*), as follows:

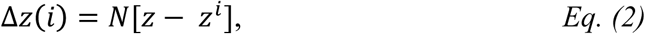

where *z* is the SC-FC correlation, *z*^*i*^ is the SC-FC correlation following removal of edge *i*, and *N* is the number of edges in the network. We performed this calculation for both preictal and ictal time periods. Since we were specifically interested in edges that statistically contribute to the increase in SC-FC correlation during seizures, we defined a measure of *contribution*, *σ*(*i*), for each edge *i* in which a structural connection exists on the increase in SC-FC correlation during seizures as follows:

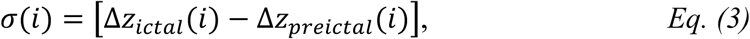

where Δz_*ictal*_ (*i*) and Δz_*preictal*_ (*i*) are the relative changes in SC-FC correlations following removal of edge *i* during the ictal and preictal periods, respectively.

We defined *contributors* of SC-FC correlation during seizures as structural edges with Δz_*ictal*_(*i*) > 0 and σ(*i*) > 0. This is because we wanted to identify regions that positively contributed to SC-FC correlation ictally, and more so ictally than preictally. To better understand the properties of contributor edges, we computed the lengths of both the contributor edges and non-contributor edges, in terms of both streamline length and physical Euclidean distance. The purpose of this analysis was to determine whether the increased SC-FC correlation during seizures was due to long- or short-range connections.

A summary of our patient-level SC-FC analysis pipeline is illustrated in **Figure 1**.

### 2.7 Statistical analyses

To determine whether the SC-FC correlations were significantly greater than chance, for each 1 second window we generated a null distribution of correlations via random permutation of the functional network edges (10,000 permutations). We then compared the mean SC-FC correlations during ictal and preictal periods with the null correlations. Next, to determine whether there was a significant increase in SC-FC correlation between preictal and ictal periods, we computed the difference between the mean ictal *z* and the mean preictal *z* for each seizure event. We modeled these paired differences using a linear mixed effects model with subject assignment as the random effect, and determined whether the difference was significantly greater than zero using the parametric bootstrap method (1000 bootstrapped samples), which is robust to small sample sizes [Davison and Hinkley, 1997; Halekoh and Højsgaard, 2014]. To assess whether the findings were robust to our choice of non-ictal period, we repeated the above analysis substituting the preictal periods with interictal periods of equivalent duration that were at least 6 hours away from seizure activity (Note: data from these interictal periods were verified via visual inspection to be artifact-free and were processed as in *2.5*). To further compare findings during preictal and interictal periods, we also carried out the above statistical analysis to determine significant differences between mean preictal *z* and mean interictal *z*. To assess the degree of intra-subject similarity of SC-FC evolution, we compared the between-subject Euclidean distances (described in *2.6*) to the within-subject Euclidean distances and tested the significance of the difference using permutational multivariate analysis of variance (PERMANOVA) (999 permutations) [Anderson, 2017].

To characterize the properties of edges that contribute to the increase in SC-FC correlation during seizures, we computed the mean length of all contributor edges and the mean length of all non-contributor edges for each subject. Edge length was computed using two metrics: mean streamline length, and Euclidean distance. We compared the mean contributor and non-contributor edge lengths using a paired *t*-test. Furthermore, to assess the relationship between edge contribution and edge length among the contributor edges, we classified contributor edges into “low”, “medium”, and “high” contribution levels for each subject using tertiles. The edge lengths in these three categories were compared using paired *t*-tests. Finally, given prior knowledge that structural connection weights decrease with Euclidean distance [Donahue et al., 2016; Kaiser and Hilgetag, 2004; Lewis et al., 2009; Rubinov et al., 2015], we repeated all analyses after removing the effect of Euclidean distance from the structural networks using linear regression.

## 3. Results

### 3.1 Clinical data

A total of 45 clinical seizures (mean duration 71 s +/- 44 s), were recorded across the 9 patients (mean age 40.2 +/- 11.8; 5 female). All seizures had focal onset, and were characterized as focal aware, focal impaired awareness, or focal to bilateral tonic-clonic. Patient demographic and clinical details are detailed in **Table 1**.

**Table 1:** Patient demographic and clinical information. Post-surgical outcome was based on Engel classification score (scale: I–IV, seizure freedom to no improvement). M: male, F: female, RTL: right temporal lobe, LTL: left temporal lobe, BTL: bilateral temporal lobe, DNET: dysembryoplastic neuroepithelial tumor, PVH: Periventricular heterotopia, FIAS: focal impaired awareness seizure, FAS: focal aware seizure, FBTC: focal to bilateral tonic-clonic, ATL: anterior temporal lobectomy.

| Subject # | Age | Gender | Outcome | Localization | Seizures recorded (#) | Treatment |
| --- | --- | --- | --- | --- | --- | --- |
| 1 | 48 | F | IA | RTL | FBTC (3) | ATL + hippocampectomy |
| 2 | 39 | M | IB | RTL/DNET | FBTC (2) | ATL + hippocampectomy |
| 3 | 45 | F | IA | LTL | FIAS (1), FBTC (3) | ATL + hippocampectomy |
| 4 | 36 | M | IB | RTL | FAS (1), FIAS (5) | ATL + hippocampectomy |
| 5 | 40 | F | IA | RTL | FIAS (5) | ATL + hippocampectomy |
| 6 | 50 | M | N/A | Bilateral PVH | FIAS (4) | N/A |
| 7 | 24 | M | 1B | RTL | FIAS (6) | ATL + hippocampectomy |
| 8 | 58 | F | N/A | BTL | FIAS (5) | N/A |
| 9 | 22 | F | 1A | LTL | FIAS (10) | Laser ablation |

### 3.2 SC-FC coupling using broadband functional connectivity

To assess the overall temporal patterns of SC-FC coupling changes during seizures, we first quantified SC-FC correlations using broadband functional connectivity networks. For each individual seizure event, we determined the degree of SC-FC coupling, as measured by *z* (**Figure 2A**). For all seizures in all subjects, SC-FC coupling was significantly greater than chance during interictal, preictal and ictal periods (*p* < 0.05, permutation-based testing; **Figure 2B**). While the temporal progression of SC-FC changes was subject-specific (**Figure 2C**), there was a consistent increase between preictal and ictal periods (**Figure 2D**). Per-seizure paired differences in mean *z* values reveal significantly greater SC-FC correlation during ictal periods than preictal periods (*p* = 0.023, linear mixed effects analysis with subject as random effect). This effect was maintained when substituting preictal periods with randomly chosen interictal clips of equivalent duration at least 6 hours away from seizure activity (*p* = 0.021). It was also maintained after regressing out the effect of distance (*p* < 0.05, **Supp. Figure 2**). Moreover, there were no significant differences between preictal and interictal period SC-FC correlation values (*p* = 0.70).

**Figure 2:**
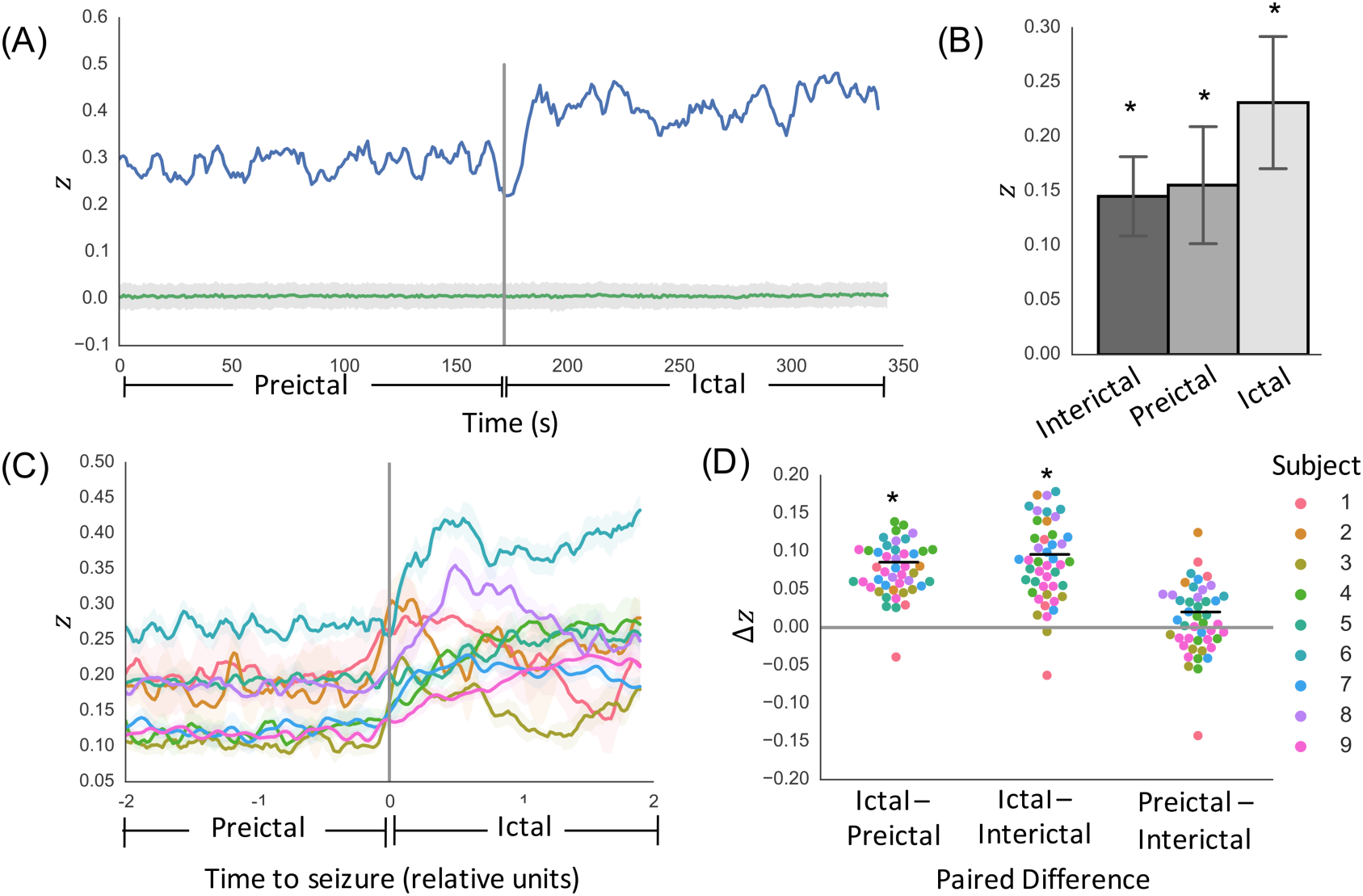
SC-FC analysis using broadband functional connectivity. **(A)** Temporal dynamics of SC-FC correlation as measured by Fisher’s *z* for one example seizure in one patient, along with permutation-based null distribution of *z* values (mean +/- standard deviation). **(B)** Per-seizure *z* values during interictal, preictal, and ictal periods reveal SC-FC correlations significantly greater than chance across all periods (*p* < 0.05). **(C)** Temporal dynamics of SC-FC correlation across all subjects (mean +/- standard deviation across seizures in each subject). For visualization purposes only, time courses were normalized to span 200 evenly spaced time windows (100 preictal and 100 ictal) and smoothed with a 5-window moving average filter. **(D)** Per-seizure paired differences in mean *z* values reveal significantly greater SC-FC correlation during ictal periods than preictal periods (*p* = 0.023). This effect holds when substituting preictal periods with interictal periods (*p* = 0.021), with no significant difference between preictal and interictal period SC-FC correlation values (*p* = 0.70).

### 3.3 Frequency-specific SC-FC analysis

Next, to better understand the frequency dependence of the observed increase in SC-FC coupling during seizures, we repeated the SC-FC coupling analysis across four frequency bands (α/θ, β, low-γ, and high-γ). Similar to the previous analysis, we found that the extent of SC-FC coupling was significantly greater than chance at all time points during preictal and ictal periods (*p* < 0.05, permutation-based testing) for all frequency bands (**Figure 3A**). Moreover, while the preictal SC-FC was lower in higher frequency bands (**Figure 3B**), the increase in SC-FC coupling between preictal and ictal periods was significant across all frequency bands (α/θ: *p* < 0.05; β: *p* < 0.05; low-γ: *p* < 0.05; high-γ: *p* < 0.05) (**Figure 3B, 3C**). This finding was upheld after regressing out the effect of distance (**Supp. Figure 2**). Similar to the findings with broadband functional connectivity, the findings were consistent when substituting preictal periods with interictal periods (α/θ: *p* < 0.05; β: *p* < 0.05; low-γ: *p* < 0.05; high-γ: *p* < 0.05), and there were no significant differences between preictal and interictal period SC-FC correlation values (α/θ: *p* < 0.05; β: *p* < 0.05; low-γ: *p* < 0.05; high-γ: *p* < 0.05) (**Supp. Figure 3**).

**Figure 3:**
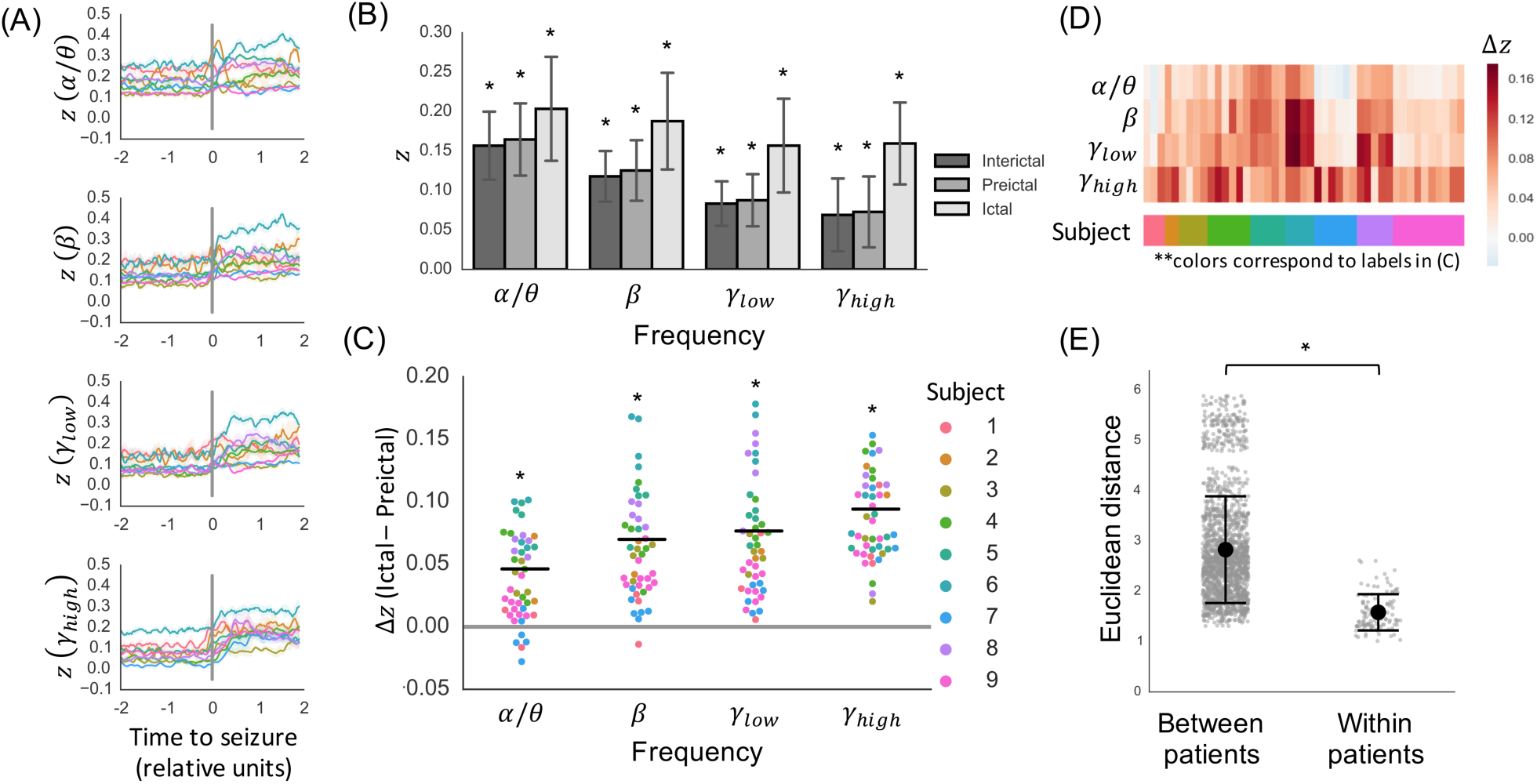
Frequency-specific SC-FC analysis. **(A)** Temporal dynamics of SC-FC correlation as measured by Fisher’s *z* in alpha/theta, beta, low gamma, and high gamma frequency bands (mean +/- standard deviation across seizures in each subject, following interpolation to normalize ictal and preictal durations). **(B)** Per-seizure *z* values during interictal, preictal, and ictal periods (mean +/- S.D.) are significantly greater than chance (*p* < 0.05, permutation-based testing). **(C)** The increase in SC-FC correlation between preictal and ictal periods is further illustrated using paired differences for each individual seizure (*p* < 0.05, linear mixed effects analysis with subject as random effect). **(D)** Heatmap illustration highlights that frequency-dependent changes in SC-FC correlation are subject-specific. **(E)** Seizures within subjects evolve similarly, as evidenced by higher between-patient Euclidean distances between SC-FC correlation time courses compared to within-patient distances (*p*<0.001, *R*^2^=0.50, permutational MANOVA).

We noted that while the increase was significant across all frequency bands, there were subject-specific frequency-dependent changes in SC-FC correlation. For example, subject 4 exhibited particularly salient increases in SC-FC_high-γ_ coupling, while subject 6 had only moderate increases in SC-FC_high-γ_ coupling but higher increases in SC-FC_β_ and SC-FC_low-γ_ (**Figure 3C**, **Supp. Figure 4**). To quantify this subject-specific effect, we determined the most salient frequency band for each subject by identifying the band with the maximum mean increase in SC-FC coupling across seizure events (**Figure 3D**).

**Figure 4:**
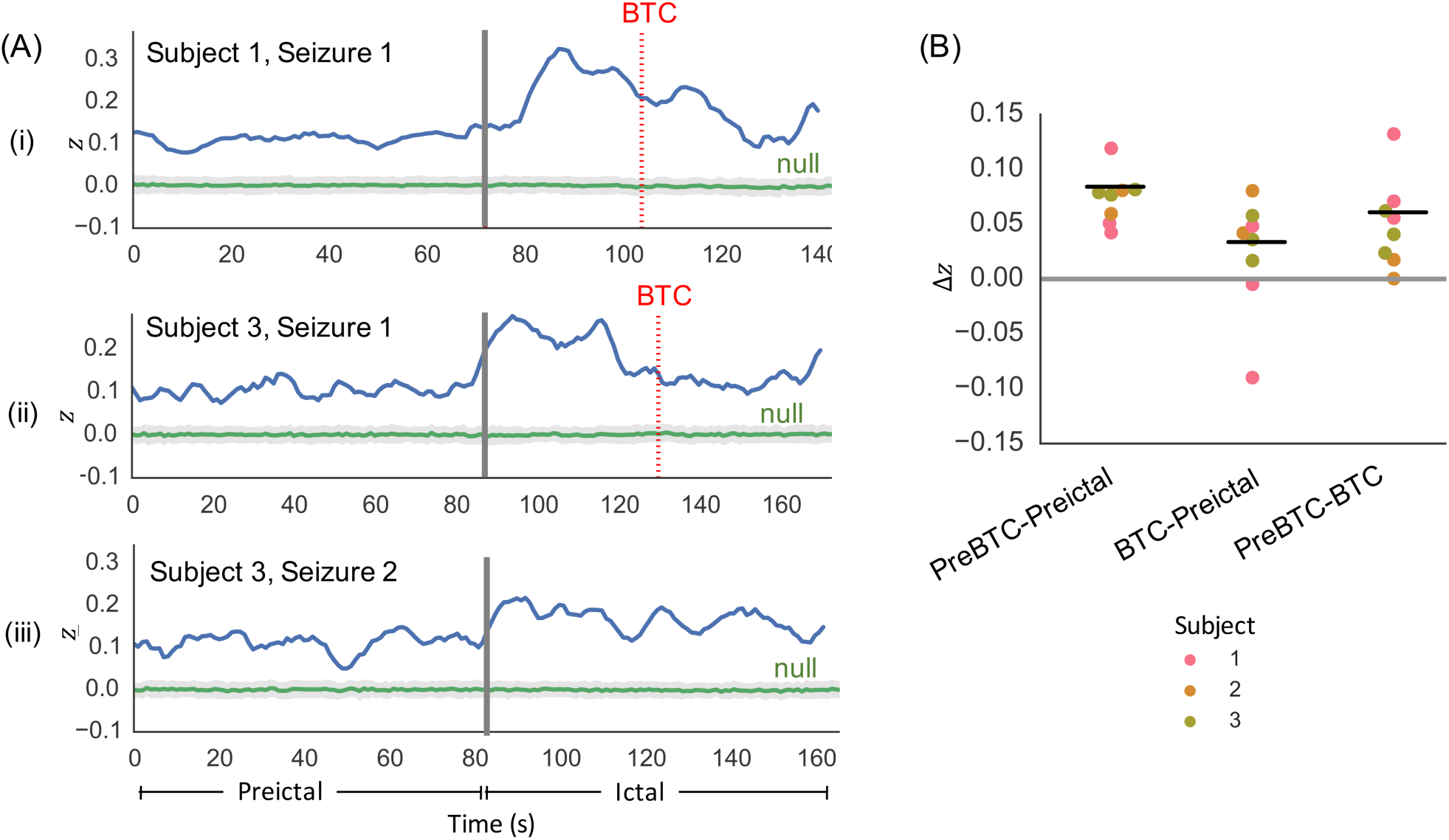
Assessment of SC-FC coupling in focal to bilateral tonic-clonic seizures. (**A**) Illustration of SC-FC coupling in two focal to bilateral tonic-clonic seizures, one from Subject 1 and one from Subject 3, reveals decrease in SC-FC coupling following bilateral tonic-clonic (BTC) onset (BTC onset indicated by dotted red line). For comparison, SC-FC coupling time course from a focal impaired awareness seizure in Subject 3 (without BTC) does not illustrate the same decrease. (**B**) In all bilateral tonic-clonic seizures, per-seizure paired differences in mean *z* values reveal significantly greater SC-FC correlation during pre-BTC ictal periods than preictal periods (*p* < 0.05), as well as significantly greater SC-FC correlation during pre-BTC ictal periods than post-BTC ictal periods (*p* < 0.05).

Finally, we characterized the within-subject similarity of the SC-FC time courses across all frequency bands. Using Euclidean distance as a measure of dissimilarity, we determined that the SC-FC time courses were significantly more similar within-patient than between-patient (*p*<0.001, *R*^2^=0.50, permutational MANOVA) (**Figure 3E**), indicating that the temporal dynamics of SC-FC coupling is stereotyped in each patient across seizure events.

### 3.3 SC-FC sub-analysis in focal to bilateral tonic-clonic seizures

As previously noted, the temporal progression of SC-FC changes was subject-specific (**Figure 2C, Figure 3A**). More specifically, we observed that in patients who experienced focal to bilateral tonic-clonic (FBTC) seizures (subjects 1-3), there was a drop in SC-FC coupling after the initial rise following seizure onset. Analysis of the individual SC-FC time courses in these seizures revealed that the drop corresponded with onset of BTC activity (**Figure 4A**). Furthermore, quantitative analysis revealed significantly greater SC-FC correlation during pre-BTC ictal periods than preictal periods (*p* < 0.05), as well as significantly greater SC-FC correlation during pre-BTC ictal periods than post-BTC ictal periods (*p* < 0.05). To focus on the relationship between structure and function prior to the onset of generalized hypersynchronous activity, we limited the ictal periods to the periods prior BTC onset for the subsequent virtual edge resection analysis.

### 3.4 Virtual edge resection analysis

Given our finding that SC-FC coupling was significantly higher during ictal periods compared with preictal periods, we wanted to identify and characterize the structural edges that statistically accounted for this increase. After quantifying the contribution *σ*(*i*) of each edge on the SC-FC coupling, we mapped the contributor edges onto each subject’s brain (**Figure 5**) to facilitate subject-specific characterization of SC-FC relationships. Furthermore, at the group level, we determined that contributor edges were predominantly short-range, as quantified by significantly shorter edge lengths in contributors compared with non-contributors, based on both geometric Euclidean distance (*p* < 0.05, two-tailed paired *t* test) (**Figure 6A**) and streamline distance (*p* < 0.05, two-tailed paired *t* test) (**Figure 6B**). This finding held individually for each subject. Furthermore, within the contributor edges, we found a trend within each subject that higher contribution edges are shorter-range, both in terms of Euclidean distance (**Figure 6C**) and streamline length (**Figure 6D**). These findings held following distance regression (**Supp. Figure 2**).

**Figure 6:**
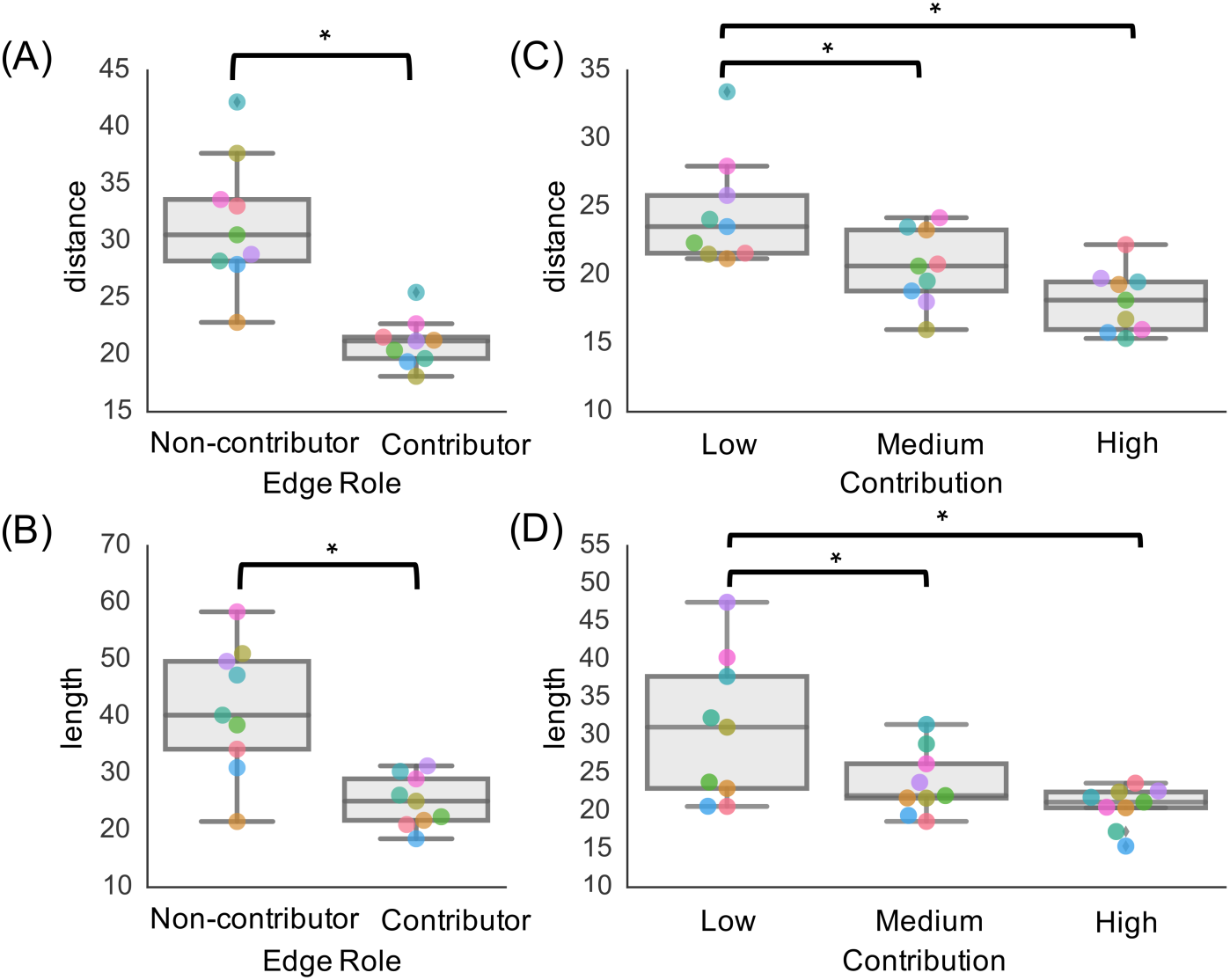
Relationship between edge contribution and edge length. Findings reveal that contributor edges are shorter-range in terms of both (**A**) Euclidean distance and (**B**) streamline length (*p* < 0.05, two-tailed paired *t* test). Furthermore, there is a trend that edges with higher contribution are shorter-range, in terms of both (**C**) Euclidean distance and (**D**) streamline length, with significant differences between low and medium contribution edges (*p* < 0.05, two-tailed paired *t* test), and low and high contribution edges (*p* < 0.05, two-tailed paired *t* test).

**Figure 5:**
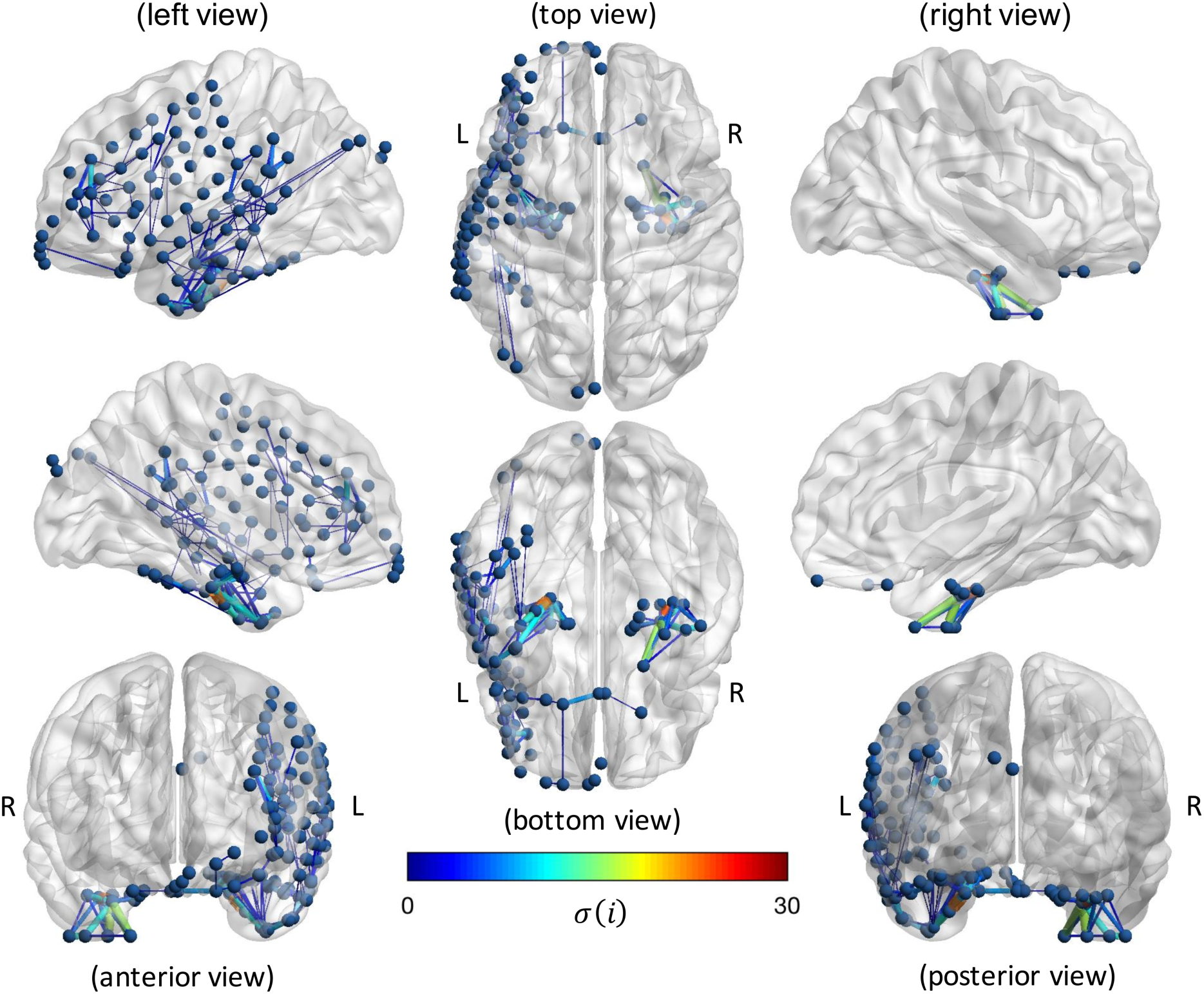
Subject-specific virtual edge resection approach to determine the contribution, *σ*(*i*), of each structural edge *i* on the increase in SC-FC correlation during seizures. Results are shown for an example seizure in a patient with left temporal lobe epilepsy. Only “contributor” edges (*σ*(*i*) > 0 and Δ*z*_*ictal*_ (*i*) > 0) are included to highlight edges that are associated with the SC-FC increase, with edge thickness and color used to representing magnitude of *σ*(*i*).

## 4. Discussion

The main goal of this study is to characterize the relationship between structural and functional connectivity during seizure onset and spread. Using network-based analysis of HARDI and iEEG data, we observe significant structure-function coupling at rest and a marked increase in this coupling during the progression from preictal to ictal states. This finding persists across frequency bands, with subject-specific levels of frequency-dependent increases. Furthermore, we present a technique for assessing the impact of individual structural connections to the observed ictal increase in structure-function correlation, and demonstrate that the effect is primarily due to short-range connections. Consistency of findings across seizures within each patient suggest that the spatiotemporal patterns of structure-function coupling are highly stereotyped. Our findings shed light on the dynamics of focal epileptic seizures in relation to underlying structure by demonstrating that seizure spread is tightly controlled by short-range structural connections.

### 4.1 Structure-function coupling across time, frequency and space

We observe greater coupling between structural and broadband functional networks during preictal and interictal periods than expected by chance. This finding is consistent with studies relating DTI-based structural networks with resting-state fMRI-based functional networks in healthy adults [Damoiseaux and Greicius, 2009; Hermundstad et al., 2013; van den Heuvel et al., 2009; Honey et al., 2009; Skudlarski et al., 2008; Zhang et al., 2010]. It is important to note that the functional signals recorded using iEEG are fundamentally different from those recorded using fMRI. While recent studies suggest that blood-oxygen level dependent (BOLD) signal fluctuations correlate with slow fluctuations in EEG gamma power, the exact relationship between fMRI (BOLD) signals and electrophysiology has yet to be resolved [He and Liu, 2008; He and Raichle, 2009; Ko et al., 2011; Logothetis et al., 2001] Nonetheless, our finding suggests that the tie between structure and function at rest is robust across diverse measurements of functional connectivity.

Interestingly, we observe that preictal FC networks in lower frequency bands have higher correlation to SC networks than preictal FC networks in higher frequency bands. This relationship decreases during the ictal period, with high SC-FC coupling in all frequency bands. Since it is believed that lower frequencies facilitate long-distance connections in the brain while higher frequencies facilitate shorter connections [Kopell et al., 2000; Miller et al., 2007], our finding may suggest a relative shift to short-range, high-frequency connectivity during seizure generation. However, this observation could be influenced by the spatial distribution of electrodes, which tend to be clustered around the putative seizure onset zone, leading to a bias towards short-range connections within the seizure generating network. Therefore, we plan to corroborate these findings in patients with stereoelectroencephalography (stereo-EEG), an increasingly popular and less invasive method that records from stereotactically placed intracranial depth electrodes and allows for wider sampling of the brain network [Cossu et al., 2005; Luders et al., 2013; Varotto et al., 2012].

Despite individual variations inherent to our patient population, the finding of increased SC-FC coupling during ictal periods compared with preictal periods is extremely robust, using both broadband and narrow-band functional connectivity. We compare SC-FC time courses from ictal periods to those of immediately preictal periods to allow for matched pairwise comparisons and to facilitate visualization along a continuous temporal scale. To ensure that activity immediately prior to seizure onset is a good representation of non-ictal activity, we repeat our analysis after substituting the preictal periods with interictal periods far away from seizure activity, and attain consistent results. The rise in SC-FC coupling during seizures indicates that seizures may rely on the brain’s underlying architecture during initial seizure spread. We note that in several of the patients, the rise in SC-FC correlation occurs prior to the clinically-marked earliest electrographic change (EEC) representative of seizure onset, suggesting that SC-FC coupling may also be a valuable biomarker for seizure prediction or its early generation.

We discover that in focal to bilateral tonic-clonic seizures, there is a significant decrease in SC-FC coupling before onset of bilateral tonic-clonic activity. This finding is not surprising, given that bilateral tonic-clonic periods are associated with generalized hypersynchronous neural activity that is not localized to particular brain regions or pathways. This finding also supports that the observed SC-FC coupling increase during seizures relates to seizure propagation, and is not simply a result of highly synchronous activity.

Of note, the temporal dynamics of SC-FC coupling is highly consistent between seizures within each patient. This indicates that seizures may be “hard-wired” in a sense, and is a macroscopic analog to the microscale finding of stereotyped ictal progression [Wenzel et al., 2017]. However, since our dataset consists of a relatively small group of adult focal epilepsy patients, with the majority having temporal lobe epilepsy, these conclusions may be specific to our dataset and should be confirmed using larger, more diverse patient populations.

To assess the role of each structural connection on the rise in SC-FC coupling during seizures, we implement a virtual edge resection method. Such leave-one-out simulation-based methods have been gaining popularity to probe the role of individual nodes and edges on overall network topology [Alstott et al., 2009; Honey and Sporns, 2008; Khambhati et al., 2016; Rafal et al., 2015]. In our case, we determine the contribution of structural connections to SC-FC correlation and generate seizure-specific brain maps of these connections. Group-level analysis reveals that connections with high contribution are predominantly short-range, in terms of both streamline length and Euclidean distance. While the connections themselves are short, the locations of these connections appear distributed across the brain, including connections that are contralateral to seizure onset. This suggests that seizure dynamics rely on a distributed network of locally clustered connections. While further analyses and validation are needed, mapping connections in relation to seizure onset and spread could ultimately be useful in pinpointing networks for therapeutic removal via targeted methods such as laser ablation [Willie et al., 2014] or neurostimulation [Fisher and Velasco, 2014].

### 4.2 Methodological Considerations and Limitations

An important but inevitable limitation of this work relates to the incomplete sampling of the network via iEEG. Electrode placement is limited by clinical necessity and constrained by the boundaries of the craniotomy, in order to minimize invasiveness and reduce patient morbidity. Therefore, it is not possible to sample functional connectivity from the entire brain at high resolution time scales accessible through iEEG. While clinicians aim to place electrodes around putative seizure onset zones, it is possible that the entire seizure network may not be captured in certain cases. Recent efforts to map whole-brain iEEG using recordings from multiple subjects [Betzel et al., 2017] and to construct models of whole-brain iEEG within individual subjects [Owen and Manning, 2017] may help circumvent this issue. Furthermore, while limited by impedance from the skull and inability to localize subcortical activity, ictal scalp EEG recordings, or ictal MEG recordings could supplement our intracranial analysis as both allow for consistent, grid-like spatial sampling with temporal resolution comparable to iEEG. The feasibility of such approaches has already been demonstrated in a recent paper revealing significant overlap between DTI networks and scalp EEG functional networks in the interictal state [Chu et al., 2015], and in early work on ictal MEG (M. Cook, S. Plummer, personal communication, 8/2018).

Our structural network findings are also limited by the capacity of our imaging methods. While HARDI has demonstrated superiority over conventional DTI in terms of its ability to resolve crossing fibers in regions of high fiber heterogeneity [Tuch et al., 2002], HARDI tractography is still only a proxy for true white matter pathways. Similar to other neuroimaging modalities, it is subject to partial volume effects and artifacts such as eddy current and susceptibility distortions [Assaf and Pasternak, 2008; Le Bihan et al., 2006]. Diffusion-based tractography is documented to recapitulate known pathways types including the short and long association fibers linking cortical gyri, the projection fiber connecting the cortex to lower portions of the brain, and the commissural fibers linking the two hemispheres [Mamata et al., 2002], but may not reconstruct unmyelinated intracortical axons. Furthermore, streamline count may not be a direct measure of the strength of anatomical connectivity.

Due to the strong relationship between spatial proximity and structural connection strength, it is not possible to entirely disentangle the effects of Euclidean distance on our findings. Given prior work that epileptiform activity propagates within layer V of the neocortex [Badawy et al., 2009], it is possible that local functional connections could partially be attributed to local cortical spreading phenomena rather than white matter propagation along short-range arcuate fibers. Local functional connectivity could also be due to measurement of a common source of signal. Despite these concerns, our finding of higher SC-FC coupling during seizures hold after regressing out Euclidean distance from our structural networks. This suggests that SC-FC coupling goes beyond solely distance-based effects.

Finally, while this study considers only direct structural connections, functional connectivity in the brain is also partially attributed to indirect structural connections [Damoiseaux and Greicius, 2009; Honey et al., 2009; Liang et al., 2017]. Future studies could employ the property of communicability [Estrada and Hatano, 2008] to incorporate path lengths of greater than one into the construction of structural networks while also accounting for the effects of spatial proximity.

### 4.3 Conclusions

We present a comprehensive approach to understanding the relationship between structure and function in the epileptic brain. Our work provides important insights into the structural underpinnings of seizure dynamics. It is our hope that by openly sharing our data and pipeline that we can accelerate translating this nascent field of network analysis in clinical epilepsy to help patients.

## 5. Acknowledgements

This work was supported by National Institutes of Health grants 1R01NS099348, K23-NS073801, 1R01NS085211, and 1R01MH112847. We also acknowledge support by the Thornton Foundation, the Mirowski Family Foundation, the ISI Foundation, the John D. and Catherine T. MacArthur Foundation, the Sloan Foundation, and the Paul Allen Foundation.

## 7. Supplementary Materials

### Supplementary Figures

**Supplementary Fig 1:**
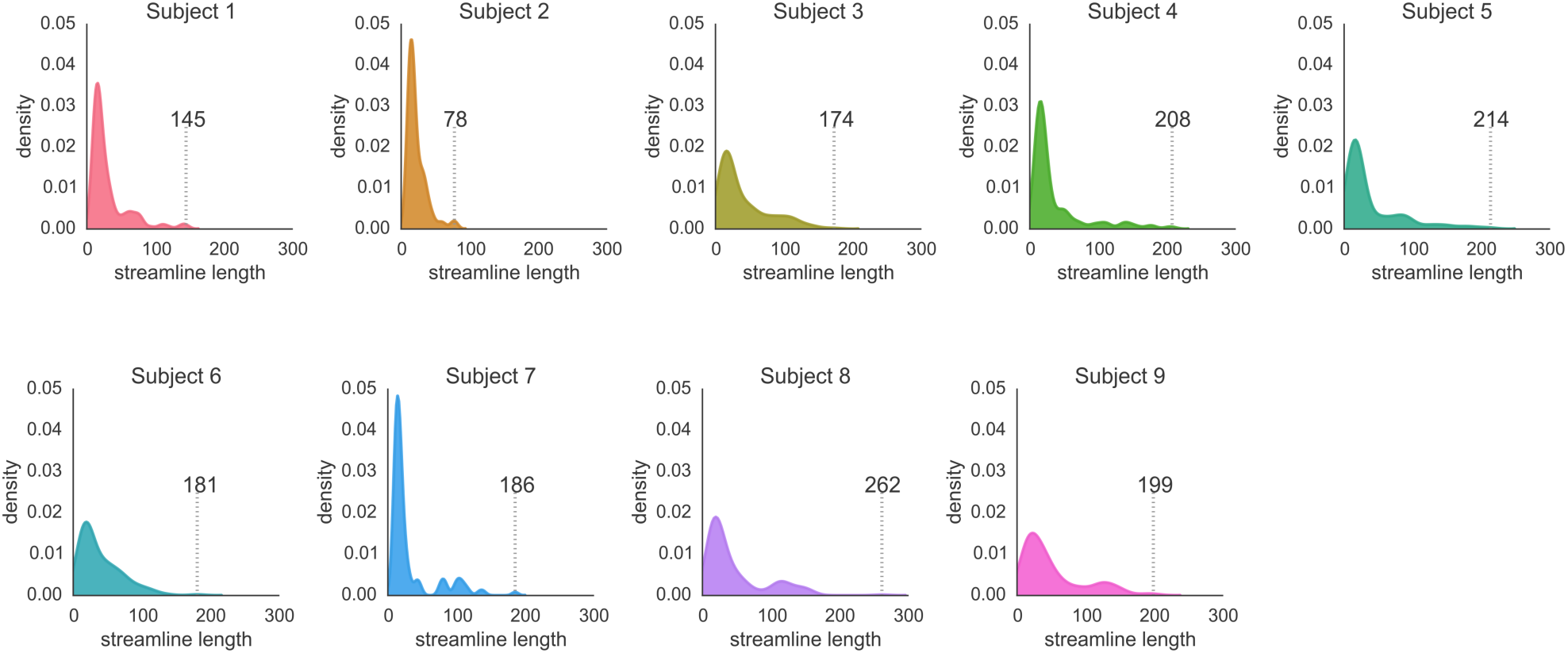
Distributions of streamline lengths for each subject, along with max streamline length for each subject (dotted grey lines).

**Supplementary Figure 2:**
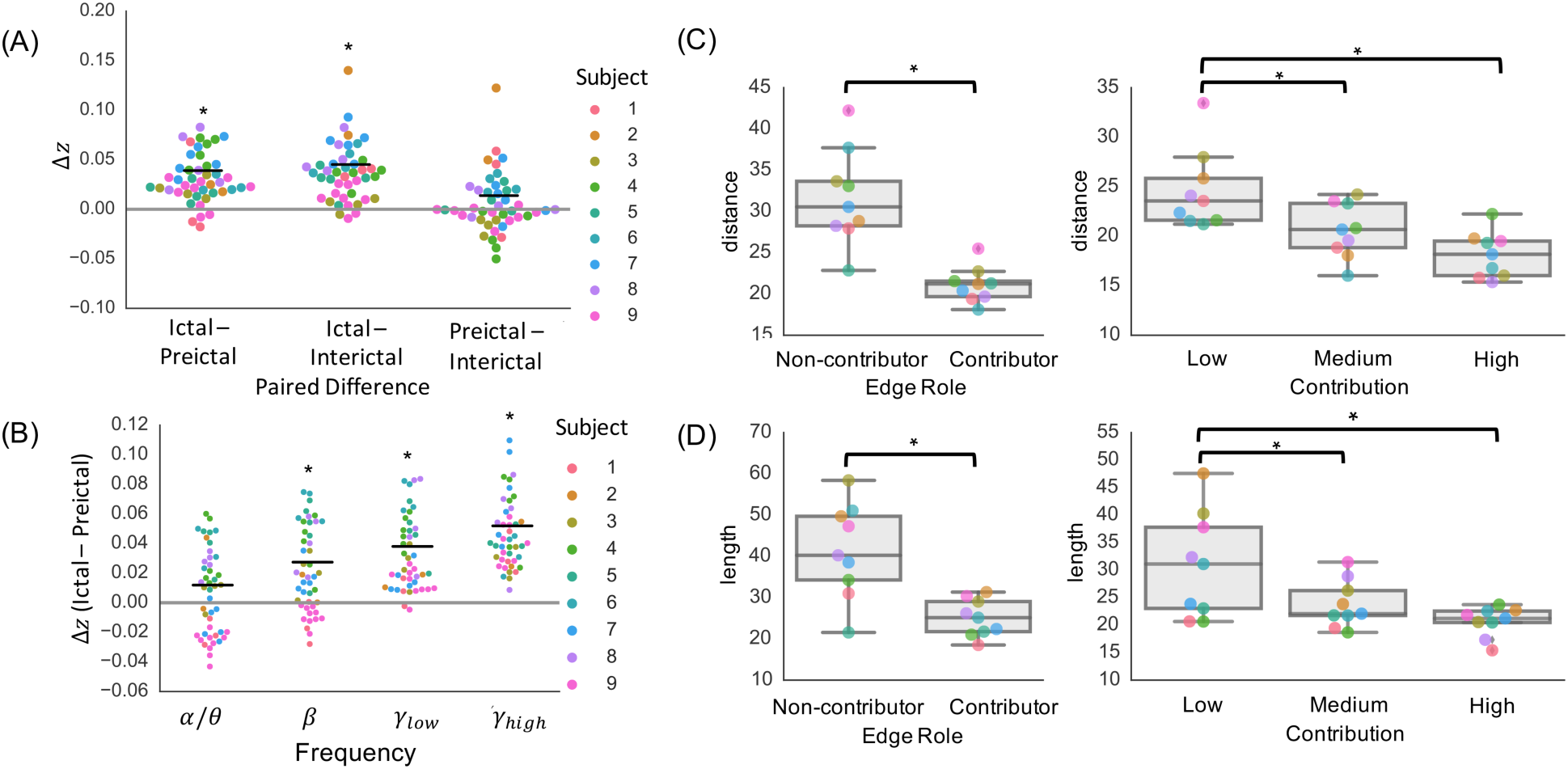
Repetition of key SC-FC coupling analyses repeated following distance regression. (A) SC-FC analysis using broadband functional connectivity. (B) Frequency-specific SC-FC analysis. (C) Relationship between edge contribution and edge length.

**Supplementary Figure 3:**
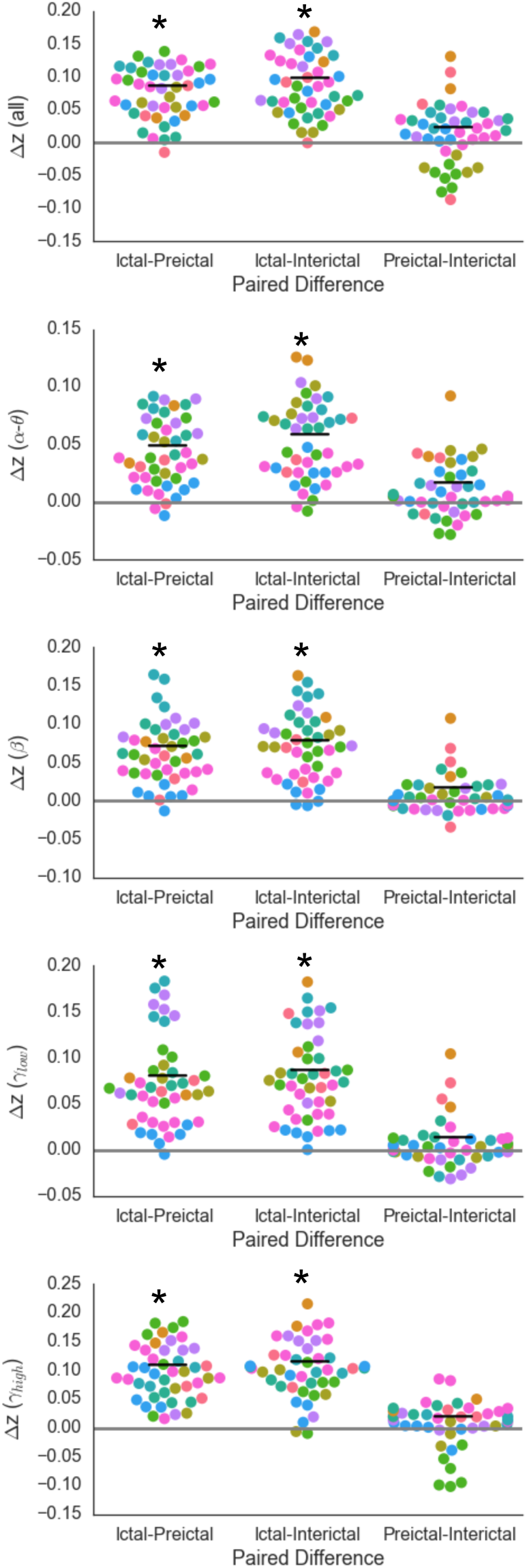
Per-seizure paired differences in mean *z* values reveal significantly greater SC-FC correlation during ictal periods than preictal periods in all frequency bands (*p* < 0.05, linear mixed effects analysis with subject as random effect). This effect holds when substituting preictal periods with randomly chosen interictal clips of equivalent duration at least 6 hours away from seizure activity (*p* < 0.05), with no significant difference between preictal and interictal period SC-FC correlation values (*p* > 0.05). See **Figures 2B** and **3C** for details.

**Supplementary Figure 4:**
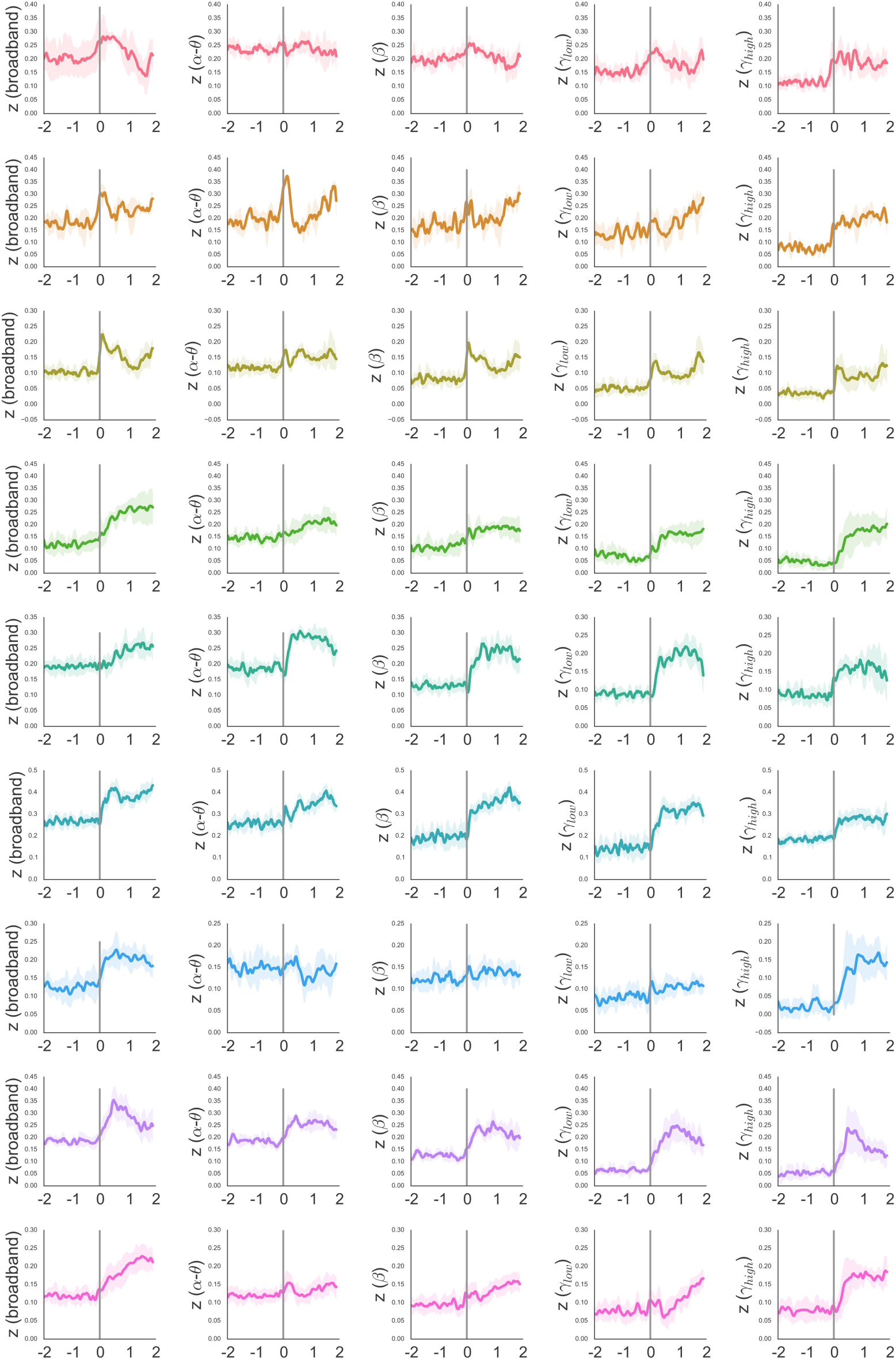
Temporal dynamics of SC-FC correlation as measured by Fisher’s *z* in alpha/theta, beta, low gamma, and high gamma frequency bands (mean +/- standard deviation across seizures in each subject, following interpolation to normalize ictal and preictal durations), delineated by subject (row/color) and band (column). See **Figure 3A** for details.

